# Sublethal chemical exposure predominantly reduces heat tolerance while increasing its variability in freshwater invertebrates and fish

**DOI:** 10.64898/2026.11.24.768878

**Authors:** Helena S. Bayat, Verena C. Schreiner, Ariane Moulinec, Ralf B. Schäfer

## Abstract

Increasing temperatures, in conjunction with widespread chemical pollution, threaten freshwater biodiversity worldwide. General patterns describing how chemical exposure affects heat tolerance have yet to be quantified beyond individual studies. Therefore we quantitatively synthesized the literature on chemical effects on the heat tolerance of freshwater fauna. We compiled heat tolerances of freshwater invertebrates and fish measured in conjunction with chemical exposure, then meta-analyzed the effect of sublethal chemical exposure on heat tolerance in terms of magnitude and variability using three-level random effects meta-analyses. We employed meta-regression to test whether these effects scale with chemical stressor intensity (acute toxicity) and differ by chemical class and taxonomic group. We found that sublethal chemical exposure consistently reduced mean heat tolerance of both fish and invertebrates with increasing toxicity across all chemical classes, with the exception of pharmaceuticals, and increased the inter-individual variation in heat tolerance measures across all chemical types. These findings imply that chemical pollution can increase freshwater species’ vulnerability to warming, while making their responses to acute warming harder to predict and affecting their ability to adapt. Thus, chemical pollution should be considered in assessments of freshwater species’ responses to ongoing climate warming.

**Summary statement:** As chemical exposure increases in toxicity, heat tolerance of freshwater fauna generally decreases, implying heightened vulnerability to heat waves in a warming world.

## 1. Introduction

Both increasing temperatures and chemical pollution endanger freshwater fauna. Freshwater ecosystems are stressed (Dudgeon, 2019) and their biodiversity is in decline (McRae et al., 2017), jeopardizing the ecological integrity of freshwaters and critical resources for human well-being (Vörösmarty et al., 2010). Heat waves are becoming increasingly frequent and severe, compounding on incremental climatic warming, with consequences for aquatic life (Barbarossa et al., 2021; Comte C Olden, 2017; Spiller et al., 2025; Tonkin et al., 2026; Xu et al., 2026). Simultaneously, over 350,000 chemicals are globally registered for use and can enter freshwater ecosystems during their production, use, and disposal (Wang et al., 2020). While effects of warming and chemicals have increasingly been investigated concurrently in the last decades, much of the research on temperature and chemicals is still conducted separately (Bayat et al., 2025; Orr et al., 2024; Polazzo et al., 2022), leaving potential interactive effects largely unexplored. Meanwhile, the majority of the extant literature on interactive effects focuses on effects of temperature on chemical toxicity rather than chemical effects on heat tolerance (Singh et al., 2025).

Acute heat tolerance, the maximum temperature that organisms can withstand as measured over short timespans in laboratory experiments, is commonly used to assess ecological consequences of warming habitats. For example, heat tolerance measures have been used to predict range shifts and risk of heat failure under various climate scenarios across taxonomic groups (Camacho et al., 2024; Jørgensen et al., 2022; Pottier et al., 2025; Verberk et al., 2020). Plasticity in heat tolerance has long been established, especially in response to acclimation temperature (Bayat et al., 2026; Pottier et al., 2022). Recently, inter-individual variation in heat tolerance estimates has received attention in the context of evolutionary adaptation to global warming (Leiva et al., 2024; Nati et al., 2021; Noble et al., 2025). Acute heat tolerance reflects multiple interacting physiological stress responses within an organism that culminate in the final endpoint (Ern et al., 2023); factors that affect the physiology of an organism can therefore change its heat tolerance. Additional stressors that affect the physiology of an organism, such as chemical pollution, potentially shift its heat tolerance and jeopardize its ability to cope with thermal challenges.

Chemical pollution has consistently been implicated as a major driver of biodiversity loss in freshwaters (Keck et al., 2025; Sayer et al., 2025). Chemicals commonly induce shifts in freshwater communities at realistic field concentrations (Beketov et al., 2013; Feckler et al., 2025; Keck et al., 2025; Liess et al., 2021; Malaj et al., 2014). Shifts in community composition could be the result of mortality, or sublethal effects on physiology that manifest in changes to growth, reproduction and homeostasis. On a physiological level, sublethal chemical effects provoke general stress responses as well as specific mechanisms (Kramer et al., 2024; Selye, 1950). General stress responses may induce protective responses towards additional stressors (Janssens C Stoks, 2017), a phenomenon termed cross-tolerance or cross-protection (Rodgers C Gomez Isaza, 2021; Sinclair et al., 2013), but may also decrease the resources available to respond to additional physiological challenges (Goussen et al., 2020; Vinebrooke et al., 2004). A specific stress response to a stressor can also confer protection to a second stressor, as is the case with the metals copper and lead (Brown, 1978). Chemicals can also induce hormesis, an effect that increases performance at low levels, which reverses at higher levels (Cedergreen, 2010; Erofeeva, 2022; Rillig et al., 2025). Organismal responses to chemicals can vary considerably between individuals of the same species. In real-world environments, inter-individual variation within a species provides the basis for selection to act upon, ultimately playing a key role in ecological community change under stress (Nakagawa et al., 2015; Violle et al., 2012). Ǫuantifying chemical effects on both the mean and variance are therefore important to understand how effects scale from the sub-organismal physiology up to the community level (McEntire et al., 2021; Schneeweiss et al., 2023).

Most studies on combined effects of warming and chemical pollution examine how temperature influences toxicity of chemicals, rather than how chemicals affect organisms’ heat tolerance (Cairns et al., 1975; Monteiro et al., 2021; Polazzo et al., 2022; Singh et al., 2025). Previous meta-analytic work has focused on the effect of temperature on the toxicity of chemicals (de Souza et al., 2023; Laskowski et al., 2010).

Work on the reverse interaction, known as “toxicant-induced climate sensitivity” or “pollutant-induced climate sensitivity”, has been developed theoretically and in individual studies, but is generally scarce (Moe et al., 2013; Noyes C Lema, 2015). To our knowledge, no studies to date have investigated how chemicals change inter-individual variation in heat tolerance, despite long-standing recognition of the biological and ecological significance of this variation (Bennett, 1987; Calow, 1996).

The impact of chemical pollutants on organism’s heat tolerance has not yet been quantified in terms of magnitude and variation across different freshwater species, despite accelerating global warming and chemical pollution. To fill this gap, we compiled heat tolerances of freshwater invertebrates and fish measured in combination with chemical exposure from the ThermoFresh database (Bayat et al., 2025), then meta-analyzed the effect of sublethal chemical exposure on heat tolerance in terms of magnitude and variation. We assessed 117 effect sizes for the change in magnitude, from 22 species exposed to 15 chemicals at various concentrations, comprising 34 distinct species-chemical combinations; for the change in variation we used 87 effect sizes, encompassing 15 species, 12 chemicals, and 18 distinct species-chemical combinations. We tested the following hypotheses with three-level random effects meta-analyses and meta-regression: (1) sublethal chemical exposure changes the mean and increases the variation of heat tolerance, (2) magnitude and variation of changes differ between organism groups (fish and invertebrates), (3) magnitude and variation of changes differ between chemical classes, and (4) changes in magnitude and variation increase with acute toxicity of the chemical exposure.

## 2. Methods

### 2.1. Data selection and preparation

Heat tolerance data tested under chemical exposure was extracted from the ThermoFresh database (Bayat et al., 2025). Data was selected based on the following criteria: (1) the dynamic method used for heat tolerance with critical thermal maximum (CTmax) or lethal thermal maximum (LTmax) as a metric (for a description of the dynamic method see work by (Bayat et al., 2026; Raby et al., 2025)), (2) waterborne chemical exposure, (3) control test conducted within the same study with identical test conditions, barring chemical exposure. Three exposures with a potential chemical toxicity of log toxic unit (logTU) smaller than -3 (calculated via Equation 1, see section 2.1.3) were excluded from subsequent analysis, as their logTU was much smaller than the rest of the data, providing insufficient information for reliable estimation below logTU of -3 (**Figures S1-S2**). The selection process resulted in 117 effect sizes from 24 studies for 34 distinct species-chemical combinations (**Figure 1**, **Table S1**). These were used for the meta-analysis of mean changes, described in section 2.1.1. In total, 87 effect sizes included a measure of error or dispersion reported in the source study, from 18 studies constituting 18 species-chemical combinations (**Table S2**), which were used to calculate the effect size used in the meta-analysis of variation, described in section 2.1.2.

**Figure 1.**
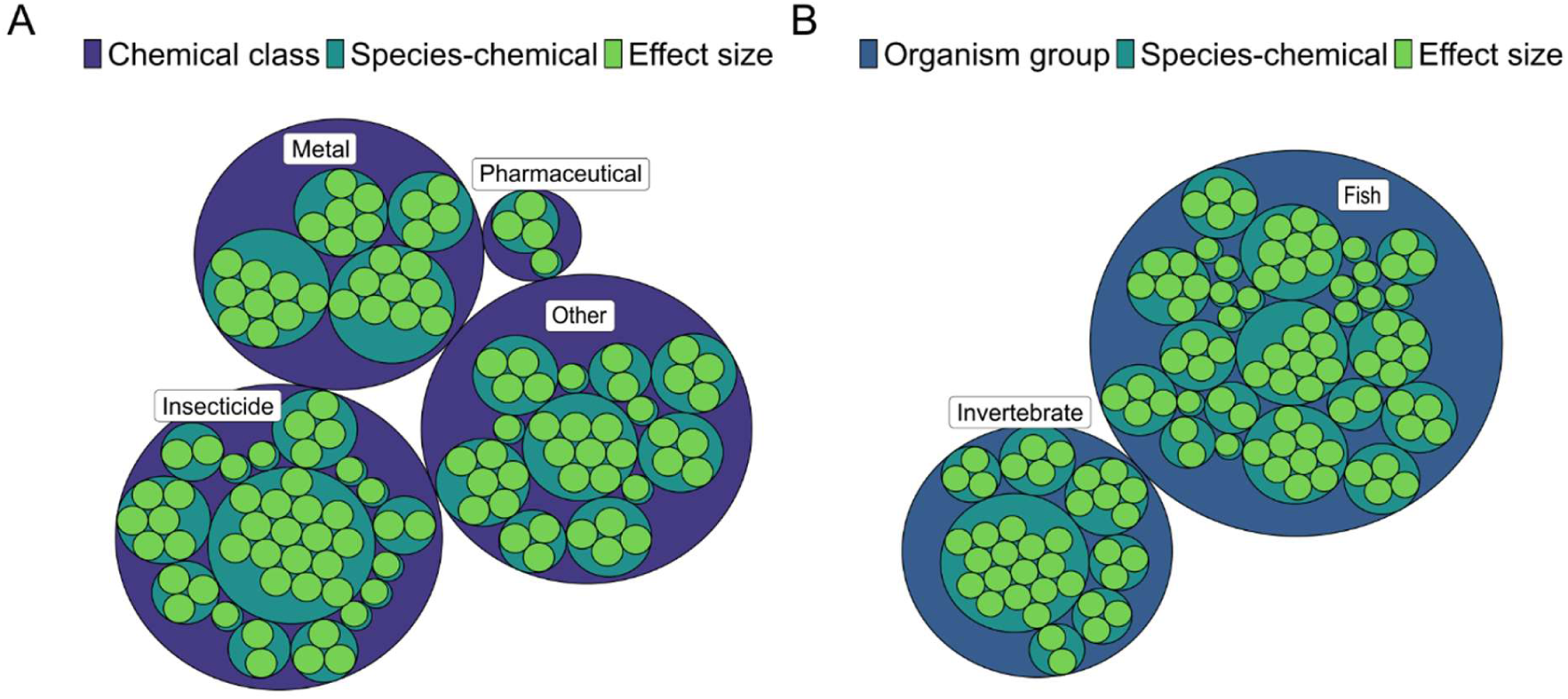
The 117 total effect sizes (mean difference in heat tolerance) analyzed in this study grouped by (A) chemical class, and (B) organism group, with species-chemical combinations as an additional grouping factor when more than one effect size was present for a given species and chemical. Each effect size (green circle) denotes a mean difference in heat tolerance under a certain set of test conditions, in this case a set of chemical exposure conditions. Multiple effect sizes within a teal species-chemical circle indicates that multiple concentrations of a chemical were tested for that species-chemical combination.

#### 2.1.1. Mean difference

Since all heat tolerances selected were determined using the same method and reported in the same units (⁰C), the mean difference between control and exposed heat tolerance was used as effect size, enhancing interpretation of the effect. This was calculated by subtracting the mean heat tolerance with chemical exposure from the mean heat tolerance for each test condition. A test condition constituted a given chemical exposure, in terms of concentration and duration, but also additional factors such as acclimation temperature. For the 26% of tests that lacked a measure of dispersion in heat tolerance, a value of dispersion was imputed. For CTmax tests under control conditions (i.e., without chemical exposure), this was the average of all CTmax standard deviations (0.662, *n* = 1138) in the ThermoFresh database, excluding those conducted under stressor exposure (both chemical stressors used in the present study, and other stressors were excluded). The average of all CTmax standard deviations in the database conducted under stressor exposure (0.809, *n* = 214) was imputed for data with chemical exposures that lacked the dispersion measure. Following imputation, the pooled variance, standard error, and 95% confidence interval limits were calculated for each mean difference following the approach recommended by Nakagawa and colleagues (Nakagawa et al., 2023). To scrutinize potential bias introduced by the dispersion imputation, we ran repeat analyses restricted to data with reported dispersion.

#### 2.1.2. Change in variation (lnCVR)

We calculated the natural logarithm of the ratio between coefficients of variation (lnCVR) between control and exposed heat tolerance as an effect size to quantify change in variation due to chemical exposure. We chose the lnCVR due to its robustness to changes in variance that scale with the mean (Nakagawa et al., 2015). The lnCVR was calculated by taking the natural logarithm of the ratio between control and exposed variance of each mean heat tolerance, for each test condition (i.e. each chemical concentration), analogous to the calculation of the mean difference. We used a second order Taylor expansion to correct for small sample size for the effect size and sampling variance calculations (Senior et al., 2020). The meta-analysis of variation was only done for the data with reported measures of dispersion (87 effect sizes, **Table S2**), as imputed values could skew the results.

#### 2.1.3. Potential chemical toxicity

The logTU, a measure of potential chemical toxicity, was used to quantify stressor intensity for each chemical exposure. We extracted both the exposure concentration and the concentration at which 50% of a test population was lethally impacted (LC_50_) for each species and chemical combination to determine logTU. The acute LC_50_ for the respective chemical, species, and test condition was extracted from the primary source study where available (for 44% of effect sizes, constituting nine species-chemical pairs), and queried from the standaRtox database if unavailable (Scharmüller et al., 2020). Wherever possible, a LC_50_ that matched the exposure duration was chosen (**Table S3**). The logTU was calculated from the LC_50_ and exposure concentration for each species-chemical combination using the following equation:

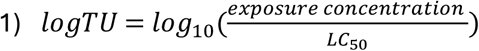

All data processing and plotting of figures was conducted in R v4.6.1 (R Core Team, 2025) using the packages dplyr v1.1.4 (Wickham et al., 2026), metafor v4.8.0 (Viechtbauer, 2026), clubSandwich v0.6.1 (Pustejovsky et al., 2026), ggplot2 v3.5.2 (Wickham, 2016), viridis v0.6.5 (Garnier et al., 2024), patchwork v1.3.2 (Pedersen, 2025), magick v2.9.0 (Ooms, 2026), cowplot v1.2.0 (Wilke, 2025), and esci v1.0.7 (Calin-Jageman, 2026).

### 2.2. Meta-analysis and meta-regression

#### 2.2.1. Model structure

Our dataset contained complex dependency structures, which are common in ecological meta-analyses (Lajeunesse et al., 2013; Nakagawa, Senior, et al., 2022; Noble et al., 2017). Chemical effects depend on both the biological target and chemical properties, therefore data from the same species or chemical can exhibit statistical dependencies. All studies in our dataset except for three (de Beeck et al., 2017, 2018; Janssens et al., 2021) tested different species-chemical combinations, which hampered the quantification of study-level dependency between effect sizes (**Table S1-S2**). Therefore, we used a three-level model, with effect sizes nested within species– chemical combinations, to account for data non-independence (**Figure 2**). As nearly every species-chemical combination originated from a separate study, this clustering was more appropriate than clustering by study. This model structure partitioned the variance into sampling error, the heterogeneity within and between each species-chemical combinations (**Figure 2**). We applied this model structure in all meta-analyses.

**Figure 2.**
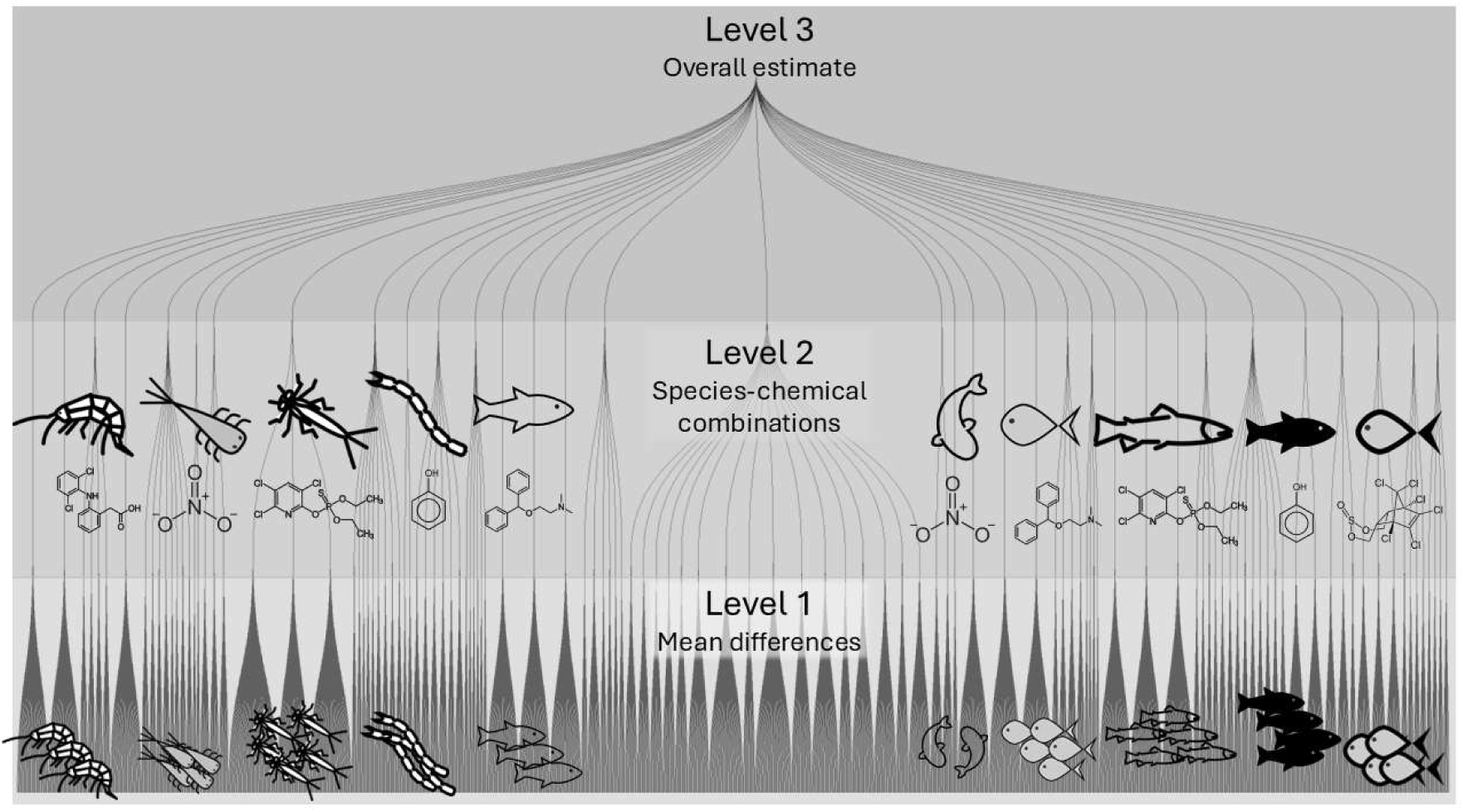
A schematic representation of the hierarchical structure of the three-level model used for meta-analyses of mean and variation in heat tolerance. The model contains three levels of aggregation to reflect dependencies within the data. Level 1 indicates the aggregation that occurs within each primary source study from individual measurements to group means per test condition (control, exposure). Heterogeneity at level 1 reflects the sampling variation. At level 2, the effect sizes are clustered according to species-chemical combinations. Nearly every species-chemical combination belonged to a separate study, therefore effect sizes were not clustered by study in our analysis. Level 3 aggregates the effect from all species-chemical clusters into one overall effect estimate.

This three-level structure allowed us to jointly test multiple concentrations for each species-chemical combination across studies, despite variations in study design. Out of 34 species-chemical combinations, 21 were tested at multiple chemical concentrations (from two up to nine), and 13 at only one concentration besides the control (green circles in **Figure 1**). As is typical for ecotoxicological tests, most studies tested multiple concentrations of the same chemical with a single control group, resulting in multiple comparisons to the control and a dependent data structure for a subset of the effect sizes that could potentially affect results (Nakagawa, Senior, et al., 2022). Therefore all models were run with an assumed correlation of *ρ* = 0.5 between effect sizes within each species-chemical grouping and robust variance estimation. (Nakagawa, Senior, et al., 2022; Nakagawa et al., 2023; Pustejovsky C Tipton, 2022; Williams et al., 2025).

We conducted meta-analyses to determine an overall effect, then subgroup analyses and meta-regression to explain heterogeneity in effect size estimates. Since we hypothesized that organism group, chemical class, and acute toxicity would explain heterogeneity in our data (hypotheses 2-4), we split our data into fish and invertebrate categories for organism group, and into insecticide, metal, pharmaceutical, and ‘other’ chemical classes (**Table S1-S2**). The ‘other’ category contained industrial chemicals and substances that have multiple uses, namely ammonia, nitrite, and selenate.

#### 2.2.2. Meta-analyses of mean difference and variation

To find out if chemical exposure altered heat tolerance and increased inter-individual variation (hypothesis 1), we meta-analyzed mean differences, calculated in section 2.1.1., and lnCVR, calculated in section 2.1.2., as effect sizes. Three-level inverse variance weighted random effects meta-analyses were run with restricted maximum likelihood estimation using the metafor (v4.8.0) package in R (Viechtbauer, 2026). We use the *I*^2^ statistic to report heterogeneity, for total heterogeneity and heterogeneity partitioned across each level of the model (**Figure 2**).

To test if chemical class, organism group, and acute toxicity explained heterogeneity (hypotheses 2-4), we conducted subgroup analyses and meta-regression for mean difference and change in variation of heat tolerance. Due to sample size constraints, we did not test all variables (organism group, chemical class, and logTU) together in the same model. First, we ran subgroup analyses separately for chemical class and organism group for both mean difference and change in variation. Then we tested logTU with chemical class and organism group, respectively, along with interactions. For the final meta-regression reported in the results, we dropped non-significant interaction terms. The variance explained by predictors was calculated according to the method developed by Nakagawa, Schielzeth, and Johnson (Nakagawa et al., 2017; Nakagawa C Schielzeth, 2013).

### 2.3. Model diagnostics, bias, and robustness

All diagnostic and robustness checks were run for the meta-analyses of mean difference in heat tolerance as well as change in variation (lnCVR). As all species-chemical combination clusters had Cook’s distance values within three times the mean Cook’s d, all data was kept for analyses. We checked for a time-lag effect, the attenuation of effect size magnitude over time, by running a meta-regression with publication year as a moderator. We assessed publication bias, wherein small studies would have larger effect sizes, by inspecting funnel plots for visible asymmetry and running an Egger regression as recommended by Nakagawa and colleagues (Nakagawa, Lagisz, et al., 2022). We conducted a sensitivity analysis for dependency with models assuming independent effect sizes, as well as correlations of *ρ* = 0.1, *ρ* = 0.5, and *ρ* = 0.9, and a model with dependency removed by selecting one effect size at the median logTU for each species-chemical combination (Lajeunesse, 2011; Noble et al., 2017). Further details are provided in the supplementary material. We report final results for models run with a *ρ* of 0.5. Data and scripts for all analyses are freely available on GitHub (https://github.com/hsbayat/sublethal_exposure_therm).

## 3. Results

### 3.1. Sublethal chemical exposure changes heat tolerance

Overall, the mean heat tolerance of fish and invertebrates decreased and inter-individual variation increased following sublethal chemical exposure. The meta-analysis of mean difference in heat tolerance resulted in an overall decrease in heat tolerance of over a degree Celsius (-1.45 ⁰C, *p* < 0.0001, **Table 1**). Of 117 total mean differences in heat tolerance, nearly 80% were negative, and 56% were negative with a confidence interval excluding zero (**Figure 3**). The meta-analysis of variation gave an overall effect estimate of 0.7, indicating an increase in inter-individual variation (**Table 1**). Of 87 total variation effect sizes, over 95% were positive, though 61% included zero in the confidence interval (**Figure 4**).

**Figure 3.**
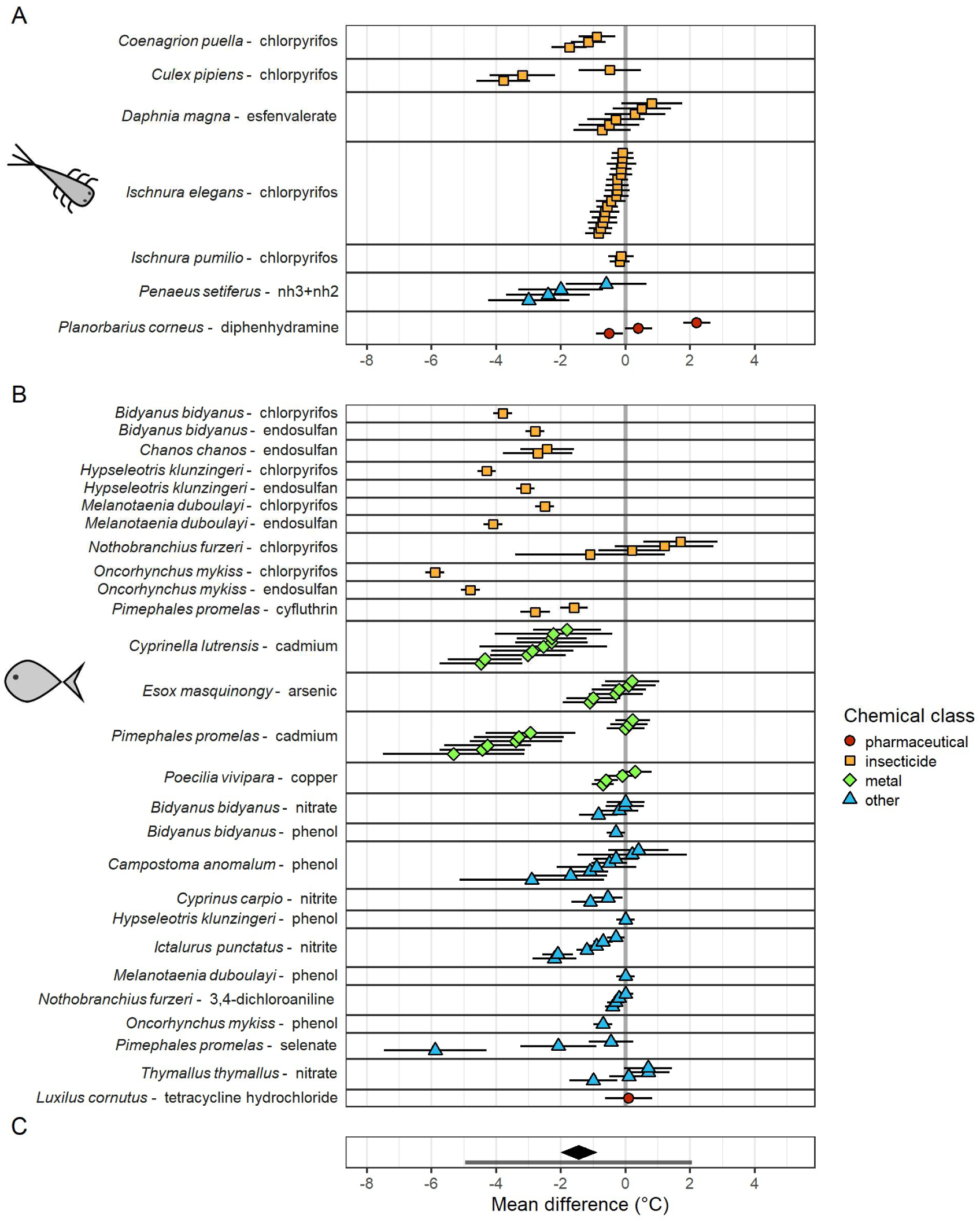
Mean difference (± standard error) in heat tolerance for (A) invertebrates and (B) fish. Multiple points per species-chemical pair indicate different chemical exposure levels. Heat tolerance effect sizes were derived from CTmax or LTmax measured with and without chemical exposure. The pooled effect (panel C) across all tests is depicted as a black diamond above the prediction interval, shown as a gray bar. The point shape and color indicate the chemical class. Negative mean difference indicates a decrease in heat tolerance, while positive mean difference indicates an increase in heat tolerance.

**Figure 4.**
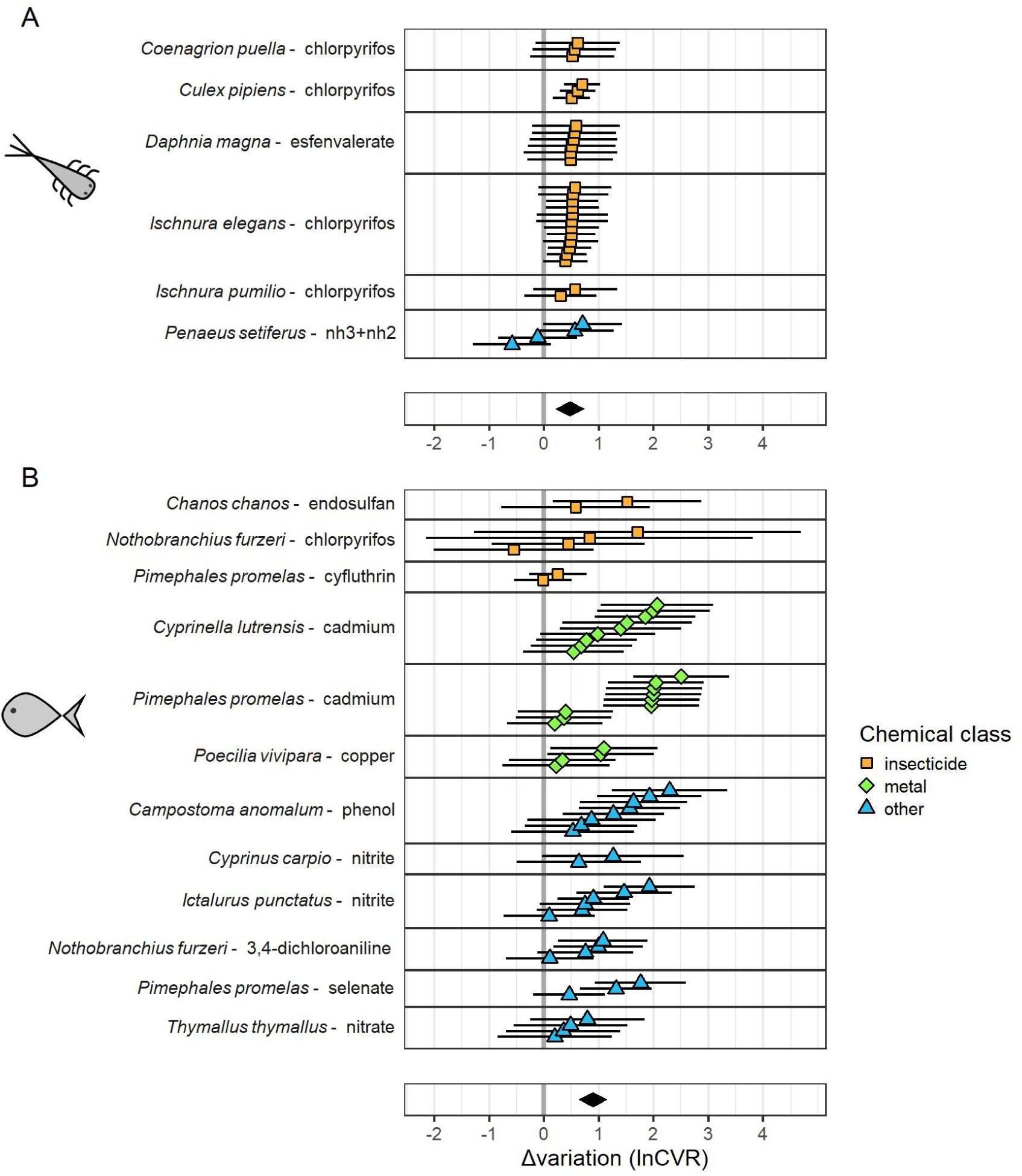
Change in inter-individual variation (± standard error) for (A) invertebrates and (B) fish. Change in variation was quantified with the logarithm of the ratios of coefficients of variation (lnCVR), where a value above and below zero denoted greater and lower inter-individual variation, respectively, in exposed organisms compared to controls. The point shape and color indicate the chemical class. The lnCVR estimate significantly differed between invertebrates and fish (*p =* 0.02). The respective overall effect diamond for each organism group is depicted as a black diamond for (A) invertebrates and (B) fish. The overall estimate for lnCVR was 0.5 for invertebrates and 0.9 for fish.

**Table 1.**
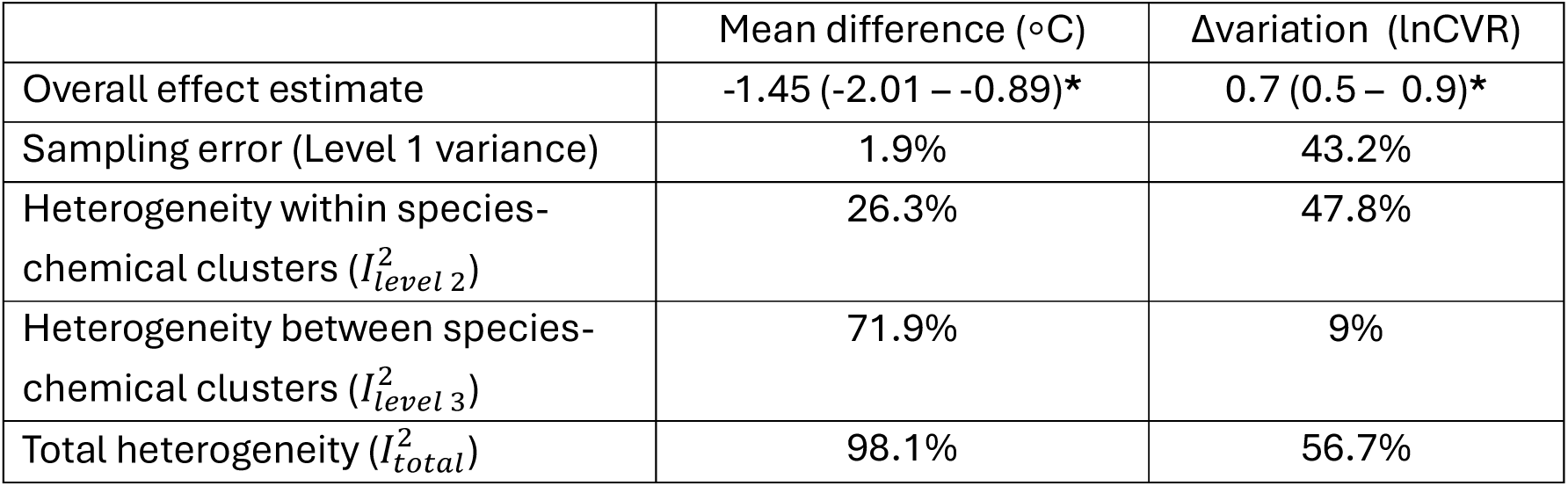
Meta-analysis results for mean difference (⁰C) and change in variation (logarithm of the ratios of coefficients of variation, lnCVR, dimensionless) effect sizes. Heterogeneity is reported as a percentage for the respective level of the model. Estimates are reported with the 95% confidence interval in parentheses. Statistically significant (*p* < 0.0001) results are marked with an asterisk (**\***).

|  | Mean difference (◦C) | Δvariation (lnCVR) |
|---|---|---|
| Overall effect estimate | -1.45 (-2.01 – -0.89)* | 0.7 (0.5 – 0.9)* |
| Sampling error (Level 1 variance) | 1.9% | 43.2% |
| Heterogeneity within species-chemical clusters ($I_{level\ 2}^{2}$) | 26.3% | 47.8% |
| Heterogeneity between species-chemical clusters ($I_{level\ 3}^{2}$) | 71.9% | 9% |
| Total heterogeneity ($I_{total}^{2}$) | 98.1% | 56.7% |

Total heterogeneity was high for the meta-analysis of mean differences in heat tolerance (*I*^2^ = 98%). The sampling variance (level 1 in the model structure, see **Figure 2**) was quite low at just under 2%. Most of the heterogeneity was present between species-chemical combinations, with an *I*^2^ of 72%, while heterogeneity within species-chemical combinations had an *I*^2^ of 26% (**Table 1**).

For the meta-analysis of variation, heterogeneity was moderate with an *I*^2^ of 57%. Heterogeneity was mostly present within species-chemical combinations (*I*^2^ = 48%), with an *I*^2^ of only 9% between species-chemical combinations. This aligns with the fact that the lnCVRs were more similar to each other than the mean differences, especially for invertebrates (**Figure 4**).

Variation of heat tolerance increased more in fish than invertebrates following chemical exposure. The overall estimate of lnCVR for fish (0.9) was nearly twice that of invertebrates (0.5), and significantly different (F(1, 10.74) = 7.01, *p* = 0.02) in the subgroup analysis (**Figure 4**). Neither chemical class (F(2, 4.9) = 4.55, *p* = 0.08) nor acute chemical toxicity (F(1, 9.2) = 1.92, *p* = 0.20), quantified by logTU, explained the increase in variation (**Figure S3**).

### 3.2. Heat tolerance decreases with increasing toxicity for all chemical classes except pharmaceuticals

Higher acute toxicity was associated with greater reductions in heat tolerance for all chemical classes except pharmaceuticals, according to the meta-regression model including chemical class and toxicity together (**Table 2**). In pharmaceuticals (*n* = 4), heat tolerance increased with toxicity, with a slope of 1.36 (**Figure 5**). The slope of logTU did not significantly differ across chemicals classed as metals, insecticides, and other types; they all had a negative relationship with logTU (slope = -0.9). The intercept for insecticides was significantly lower than for chemicals classed as other (**Table 2**). We found no significant effect of organism group on mean difference in heat tolerance, neither as sole moderator in a subgroup analysis (F(1, 9.67) = 2.55, *p* = 0.14) nor in interaction with acute toxicity (F(3, 4.5) = 3.50, *p* = 0.12).

**Figure 5.**
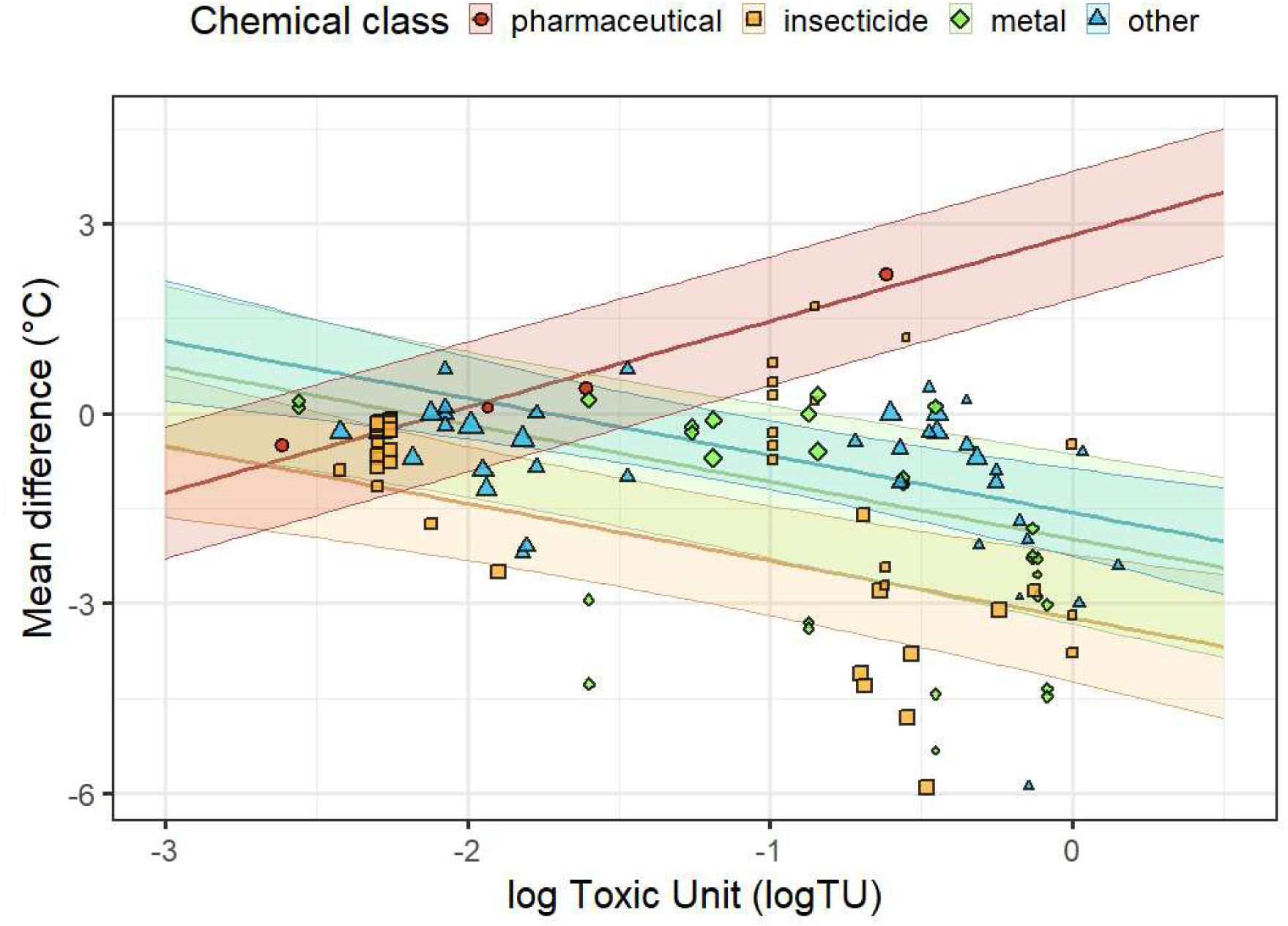
Meta-regression model results for the effect of acute toxicity and chemical class on the mean difference in heat tolerance. Points are the mean difference effect sizes for change in heat tolerance, plotted against the logTU of the chemical exposure. Point color and fill indicate chemical class, point size indicates weighting according to inverse variance. A significant interaction was found only between pharmaceuticals and logTU. Lines are plotted for each chemical class in the matching color, with confidence intervals shaded in a lighter tone, bounded by the thin lines.

**Table 2.** Subgroup analysis and meta-regression results for mean difference (⁰C) and change in variation (logarithm of the ratios of coefficients of variation, lnCVR) effect sizes. The row in bold indicates the full model with all terms, wherein ‘chemical class’ includes the terms insecticide, metal, other, and pharmaceutical for model 1, and ‘organism group’ includes invertebrate and fish terms for model 2. The subsequent rows list separate terms for each respective model. Toxicity is quantified as log toxic unit (logTU). Nonsignificant interaction terms were dropped from models reported here.

| Model | Effect size | Term(s) | Estimate | SE | P-value | Explained variance ($R_{*}^{2}$ ) |
|---|---|---|---|---|---|---|
| **1 (full)** | **Mean difference** | **Chemical class + logTU + logTU x Pharmaceutical** | **-** | **-** | **0.0001**<sup>a</sup> | $\boldsymbol{R}^{2}_{\mathrm{level2}}$**= 21.3%**<br>$\boldsymbol{R}^{2}_{\mathrm{level3}}$**= 43.8%**<br>$\boldsymbol{R}^{2}_{\mathrm{marginal}}$**= 24.8%** |
| 1 | Mean difference | Insecticide | -3.23 | 0.47 | <0.0001 | - |
| 1 | Mean difference | Metal | 1.25 | 0.56 | 0.076 | - |
| 1 | Mean difference | Other | 1.66 | 0.47 | 0.0018 | - |
| 1 | Mean difference | Pharmaceutical | 6.05 | 0.48 | 0.02 | - |
| 1 | Mean difference | logTU | -0.9 | 0.19 | 0.0012 | - |
| 1 | Mean difference | logTU x Pharmaceutical | 2.26 | 0.19 | <0.0001 | - |
| **2 (full)** | **lnCVR** | **Organism group** | **-** | **-** | **0.023**<sup>a</sup> | - |
| 2 | lnCVR | Invertebrate | 0.47 | 0.16 | 0.023 | - |
| 2 | lnCVR | Fish | 0.89 | 0.15 | 0.0002 | - |

<sup>a</sup> The *p*-value for the test of moderators of the final model, including all terms.

The final meta-regression with logTU, chemical class, and the interaction between logTU and pharmaceuticals explained 21% of the heterogeneity, as quantified by the conditional 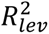, within species-chemical clusters and 44% of the heterogeneity between them, as quantified by the conditional 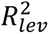 in the model (**Table 2**). The conditional R-squared values at level two and three refer to the structure of the model (**Figure 2**). This suggests that most of the heterogeneity within species-chemical estimates remains unexplained, either due to unevaluated variables or biological stochasticity. Since toxicity differed more between species-chemical clusters than within them (the range of logTU tested was more variable between studies and therefore species-chemical combinations), toxicity possessed more explanatory power in the model at the level between clusters. The final meta-regression including chemical class, logTU, and the interaction (model 1 in the first row of **Table 2**) explained more heterogeneity than logTU alone; the full model had a 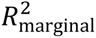 of 25% as opposed to just under 10% for a model with logTU as the sole moderator.

### 3.3. Bias and robustness

We found no concrete evidence of publication bias, nor of a shift in effects over time (time-lag effect). The funnel plots appeared symmetric upon visual inspection (**Figures S4-S5**). Neither the Egger regression nor the regression with publication year had a significant slope term (**Figure S6-S7**). The sensitivity analysis with *ρ* confirmed that dependency did not majorly shift results (**Table S4**). More details are given in the Supporting Information.

## 4. Discussion

We found that chemical exposure predominantly reduces mean heat tolerance and increases inter-individual variation therein (hypothesis 1) using meta-analyses of mean difference and change in variation. To our knowledge, no previous work has meta-analyzed the effect of chemical exposure on heat tolerance in terms of magnitude and variation in freshwater fish and invertebrates. We found high heterogeneity, as is common in ecological meta-analyses (Senior et al., 2016). The partitioning of heterogeneity indicated that mean differences in heat tolerance vary considerably depending on the species-chemical pair. We found evidence that the effects on variation but not magnitude differed by organism group (hypothesis 2), though this result should be interpreted with caution, since only one chemical was tested in both fish and invertebrates. We found a negative relationship between acute potential toxicity and change in mean heat tolerance (hypotheses 3-4) for all chemical classes, except pharmaceuticals. However, neither toxicity nor chemical class affected the variation in heat tolerance. Overall, the variables we chose to investigate (organism group, chemical class, and acute toxicity) affected mean difference and change in variation differently.

### 4.1. Ecological implications of heat tolerance reduction

The heat tolerance of fish and invertebrates decreased following sublethal chemical exposure, for all chemical classes except pharmaceuticals, implying that a multitude of freshwater species may be physiologically impaired by levels of chemical pollution commonly found in the real-world. The acute chemical exposures in this study ranged in toxicity from 0.2 to -3 logTU, representing a range of realistic chemical stressor intensities found in freshwater habitats. Field studies in Europe, North America, and Australia have measured field concentrations up to over zero acute logTU for invertebrates (Covert et al., 2020; Malaj et al., 2014; Schäfer et al., 2012; Schreiner et al., 2021; Vormeier et al., 2023), and over -1 acute logTU for fish (Cojoc et al., 2024; Covert et al., 2020; de Anda et al., 2024; Shoda et al., 2016). This means that freshwater organisms in many areas of the world regularly encounter chemical exposure levels that can provoke changes in physiological traits, compromising their ability to cope with heat waves.

In landscapes where freshwater organisms increasingly need to cope with high temperatures, a reduction in heat tolerance can have wide-ranging implications for long-term survival. The reduction in water volume associated with concurrent drought and heatwaves increases both chemical concentrations and water temperature simultaneously (Graham et al., 2024). Under these conditions, freshwater organisms face a reduction in heat tolerance when they need it most. Heatwaves, droughts, and their combination already impact freshwater biodiversity negatively (Eichelmann et al., 2025), and pollution potentially further exacerbates these negative effects. As the climate continues to warm, extremes may become the norm (Xu et al., 2026) rather than the exception for many freshwater habitats.

Heat tolerance reduction reflects physiological impairment that can have knock-on effects for coping with other adverse environmental conditions, even for species not directly affected by heat waves. A reduction in heat tolerance of 1 ⁰C for a species with a large thermal safety margin (the difference between maximum experienced environmental temperature and upper tolerance limit) will not bring that species into thermal danger. For example, certain dragonfly and aquatic beetle species have heat tolerances exceeding 40 ⁰C, while residing in temperate habitats that do not reach such temperatures in the current climate (Bayat et al., 2025; Martin et al., 1976; Sánchez-Fernández et al., 2012). A measured reduction in tolerance induced by chemicals still reflects physiological alterations that can impair organismal function. For example, zinc reduces heat tolerance by decreasing hemoglobin content (Dong et al., 2026), the same mechanism by which zinc reduces growth and reproduction in *Daphnia magna* (Vandegehuchte et al., 2010). Reduction in heat tolerance can therefore act as a biomarker for energetic and physiological costs imposed by chemicals. However, the exact mechanisms underlying heat tolerance can differ between species and type of thermal exposure (acute, chronic, fluctuating, etc) (Ern et al., 2023; Ignatz et al., 2023; Zanuzzo et al., 2019), so an effect on heat tolerance cannot directly be used to estimate effects on growth or reproduction. Future work combining theoretical bioenergetic frameworks with physiological measurements may be able to link such effects to universal patterns that exist across species (Goussen et al., 2020; Sokolova, 2021).

### 4.2. Pharmaceuticals increase heat tolerance: exceptions to the rule

While most chemical exposures led to decreases in heat tolerance in our analysis, pharmaceutical exposures presented a notable exception. Mean heat tolerance increased with increasing toxicity for pharmaceuticals (**Figure 5**). However, only two species-chemical combinations contained pharmaceuticals in our dataset; one concentration of tetracycline hydrochloride (an antibiotic) was tested in the fish *Luxilus cornutus*, with no significant effect on heat tolerance (Schubauer et al., 1980), while the antihistamine diphenhydramine resulted in increased heat tolerance in the snail *Planorbarius corneus*, driving the divergence from the other chemical classes in the meta-analysis (Kuehrer, 2017). Neither of the studies on pharmaceuticals reported measures of dispersion, so pharmaceutical effects were not included in the meta-analysis of variation. The diverging trend in pharmaceuticals we found provides only preliminary evidence that remains to be supported by follow-up studies.

This trend could be due to a compound-specific characteristic, or characteristics of the species, since both the fish *Luxilus cornutus* and the snail *Planorbarius corneus* each only appeared once in the dataset. The positive trend seen with pharmaceuticals in our study could be caused by hormesis (Cedergreen, 2010; Rix et al., 2022), and reverse at higher concentrations than tested in the original study, or be related to the substance’s mode of action. The effect on snails’ heat tolerance via diphenhydramine may have been a result of a slower behavioral response; pharmaceuticals have been found to slow response times in aquatic snails, including in the study used in our meta-analysis, and diphenhydramine is known to have sedative effects (Fong et al., 2017, 2019; Khan C Tobias, 2025; Kuehrer, 2017). Pharmaceuticals can also decrease swimming activity in freshwater crustaceans (De Lange et al., 2006; Ford C Fong, 2016), or have both stimulatory effects and inhibitory effects at different concentrations (Nielsen C Roslev, 2018; Renault et al., 2026). Running a lethal assay with the LTmax as an endpoint in addition to the CTmax would separate behavioral effects from other physiological effects caused by chemical exposure. Pharmaceuticals, or any other substances that affect behavior, may obscure physiological effects in a behavioral assay like the critical thermal method, warranting careful choice of endpoints when assessing their effects on ecologically relevant traits like heat tolerance.

### 4.3. Implications of changes in inter-individual variation

We found that chemical exposure increased inter-individual variation in heat tolerance, irrespective of toxicity level or chemical class (**Figure 4**). Inter-individual variation has been found to increase with higher temperatures in a recent meta-analysis of physiological rates (Noble et al., 2025). In our study, inter-individual variation of heat tolerance increased with chemical exposure, even at low toxicities that did not cause a notable reduction in the mean. This suggests that effects on variation can occur at lower toxicity levels than those that induce effects on mean responses. Thus many freshwater organisms are likely experiencing chemical-induced increases in variability of physiological traits, as chemical pollution is ubiquitous across the globe (Tang et al., 2021; van Vliet et al., 2023; Wolfram et al., 2026).

Not all measured variation can be attributed to ‘true’ biological variation; some variation is inherently due to sampling, handling, and experimental error. Our findings on increased variation in heat tolerance occurred nearly universally across species, chemicals, and study designs, suggesting that this variation had a true biological cause. We could not, however, pinpoint which factors drove this increase in variation. The increase in variation was not toxicity-dependent, but did differ between invertebrates and fish. The higher change in variability in fish could be due to their more complex physiology as compared to invertebrates (Noble et al., 2025), but our dataset was also biased towards fish, with nine fish and six invertebrate species, and twelve species– chemical combinations for fish opposed to six for invertebrates. Thus, the higher variation might stem from differences among species and chemicals rather than from intrinsic biological characteristics of fish or invertebrates. Additionally, none of the chemicals tested in invertebrates were tested in fish in the dataset used for the meta-analysis of variation. Metals were only tested in fish, and insecticides primarily in invertebrates. Thus, the effect of organism group could not fully be separated from the effect of the chemical class. Future work should continue to synthesize and collate new evidence to understand the causes of variation in physiological traits such as heat tolerance.

Changes in variation have consequences on the ability of populations to adapt to stress. Inter-individual variation within a population provides the basis upon which evolutionary mechanisms act, ultimately affecting how communities composed of those populations change. Standing variation in a population allows adaptations to occur much more quickly than variation via mutations, and is therefore especially pertinent in the context of rapid anthropogenic change (Barrett C Schluter, 2008). Intraspecific variability, when higher than interspecific variability, can reduce extinction rates in populations exposed to a single stressor (De Laender et al., 2014). Levels of response diversity can change whether two stressors have a synergistic or antagonistic effect on species coexistence in a plant community (Laloux C De Laender, 2026). Under multiple stressor exposure, adaptation may invoke trade-offs that can lead to increased sensitivity to later exposure combinations (Orr et al., 2022). The role of inter-individual variation in shaping trade-offs to multiple stressors remains to be thoroughly elucidated. Nonetheless, the amount of inter-individual variation within a population can determine which populations adapt and which decline under anthropogenic change.

### 4.4. Future work should consider chemical mixtures

In the real world, chemicals occur in complex mixtures (Bub et al., 2025; Finckh et al., 2024). The studies included in our analysis investigated the effect of a single chemical on heat tolerance. The pattern we found in pharmaceuticals may change with co-exposure to additional chemicals of other classes. Chemical mixtures, composed of various chemical classes, should be combined in future studies on heat tolerance to better understand effects, especially for chemical combinations that have diverging effects individually, such as the pharmaceutical diphenhydramine and the insecticide chlorpyrifos. Chemical mixtures have received considerable attention in ecotoxicology (Escher et al., 2020; Schneeweiss et al., 2023), but their effects on heat tolerance have yet to be investigated. The overwhelming diversity of chemical stressors prohibits the comprehensive investigation of all stressor combinations. However, acute heat tolerance assays offer an efficient way to screen for stressor effects (Monteiro et al., 2021), as they can be conducted rapidly and efficiently in comparison to standard ecotoxicology tests, taking a few hours as opposed to a few days.

## 5. Conclusions

Sublethal chemical exposure, with the exception of pharmaceuticals, reduced heat tolerance in both fish and invertebrates while simultaneously increasing the inter-individual variation in this trait. Therefore organisms inhabiting polluted environments face heightened susceptibility to increasingly frequent extreme events, particularly those already living near their thermal limits, as baseline temperatures increase. Accelerating anthropogenic changes present challenges to freshwater biodiversity on multiple fronts as extreme climatic events look to become the norm across the globe and chemical pollution continues. As pollution amplifies inter-individual variation in heat tolerance, efforts to predict freshwater organisms’ sensitivity to heat should explicitly incorporate this variation, especially when considering potential for adaptation. The chemicals in this study compromise only a sliver of the tremendous diversity of chemicals in use worldwide, underscoring the need for more studies encompassing a variety of stressor intensities and different chemical types, as well as chemical mixtures.

## List of symbols and abbreviations

CTmax: critical thermal maximum
lnCVR: natural logarithm of the ratio between coefficients of variation
logTU: log toxic unit
LTmax: lethal thermal maximum

## Acknowledgements

We thank all the authors of the original works that made this meta-analysis possible.

## Author contributions

**Helena S. Bayat**: Writing – Original Draft Preparation; Conceptualization; Data Curation; Formal Analysis. **Verena C. Schreiner**: Writing – Review C Editing. **Ariane Moulinec**: Writing – Review C Editing; Validation. **Ralf B. Schäfer**: Writing – Review C Editing; Conceptualization; Funding Acquisition.

## Data availability

The source data for this study are freely available on Zenodo (DOI: 10.5281/zenodo.14056759) and GitHub (https://github.com/hsbayat/ThermoFresh). All scripts and processed data will be made publicly available upon publication at following GitHub repository: https://github.com/hsbayat/sublethal_exposure_therm.

