## Supplementary Material for "Sublethal chemical exposure predominantly reduces heat tolerance while increasing its variability in freshwater invertebrates and fish"

**Supporting Information**.

Supplement to the paper: Sublethal chemical exposure predominantly reduces heat tolerance while increasing variation thereof in freshwater invertebrates and fish

**Authors**: Helena S. Bayat, Verena C. Schreiner, Ariane Moulinec, Ralf B. Schäfer

**Supplementary Tables**.

**Table S1**. Data for mean difference in heat tolerance following the selection process. Measures of dispersion were imputed for 30 mean difference effect sizes.

| Category | Subgroups | Studies | Species | Species-chemical combination | Effect sizes |
| --- | --- | --- | --- | --- | --- |
| Organism group | Fish | 15 | 15 | 27 | 80 |
|  | Invertebrate | 9 | 7 | 7 | 37 |
| Chemical class | Pharmaceutical | 3 | 2 | 2 | 4 |
|  | Insecticide | 11 | 12 | 16 | 46 |
|  | Metal | 3 | 4 | 4 | 28 |
|  | Other | 9 | 11 | 12 | 39 |
| Total | All | 24 | 22 | 34 | 117 |

**Table S2**. Data subset with reported measures of dispersion, used in the meta-analysis of variation.

| Category | Subgroups | Studies | Species | Species-chemical combination | Effect sizes |
| --- | --- | --- | --- | --- | --- |
| Organism group | Fish | 11 | 9 | 12 | 57 |
|  | Invertebrate | 7 | 6 | 6 | 30 |
| Chemical class | Pharmaceutical | 0 | 0 | 0 | 0 |
|  | Insecticide | 9 | 8 | 8 | 34 |
|  | Metal | 2 | 3 | 3 | 22 |
|  | Other | 7 | 7 | 7 | 31 |
| Total | All | 18 | 15 | 18 | 87 |

**Table S3**. The degree to which the selected LC_50_ matched the chemical exposure conditions for the log toxic unit (logTU) calculation. An exact match used a LC_50_ from the same study, species, and exposure duration as the tested exposure. When a LC_50_ from the same species was not available, a closely related organism was chosen.

| logTU match | Species-chemical combinations | Effect sizes |
| --- | --- | --- |
| Exact match (study, species, exposure duration) | 4 | 18 |
| Study, species | 9 | 33 |
| Species, exposure duration | 2 | 12 |
| Species | 7 | 23 |
| Genus | 3 | 7 |
| Family level or higher | 9 | 24 |

**Table S4**. Comparison of meta-analyses of mean difference using a three-level model assuming independent effect sizes, correlated effect sizes for *ρ* = 0.1, *ρ* = 0.5, and *ρ* = 0.9, as well as trimmed to one effect size per species-chemical grouping (in this case, the median acute chemical toxicity). All results remain significant, with minor shifts in the overall estimate and confidence intervals when effect sizes are correlated. Heterogeneity, represented by *I^2^*, was high for all models. Estimation of variance components shifted between level 2 and level 3 between the models, but level 3 variance (between species-chemical combination) remained substantially larger than level 2 variance (within each species-chemical combination) for all models run.

|  | Independent | *ρ* = 0.1 | *ρ* = 0.5 | *ρ* = 0.9 | Median |
| --- | --- | --- | --- | --- | --- |
| Estimate | -1.49 | -1.48 | -1.45 | -1.42 | -1.5 |
| Standard error | 0.28 | 0.28 | 0.28 | 0.28 | 0.3 |
| 95% CI | -2.05 – -0.93 | -2.04 – -0.92 | -2.01 – -0.89 | -1.98 – -0.85 | -2.12 – -0.88 |
| *p-*value | <0.0001 | <0.0001 | <0.0001 | <0.0001 | <0.0001 |
| Level 2 variance ($I^{2}$) | 25.06% | 25.11% | 26.25% | 29.17% | - |
| Level 3 variance ($I^{2}$) | 73.07% | 73.02% | 71.88% | 68.98% | - |
| Total $I^{2}$ | 98.13% | 98.13% | 98.13% | 98.15% | 98.76% |

**Raw data**.

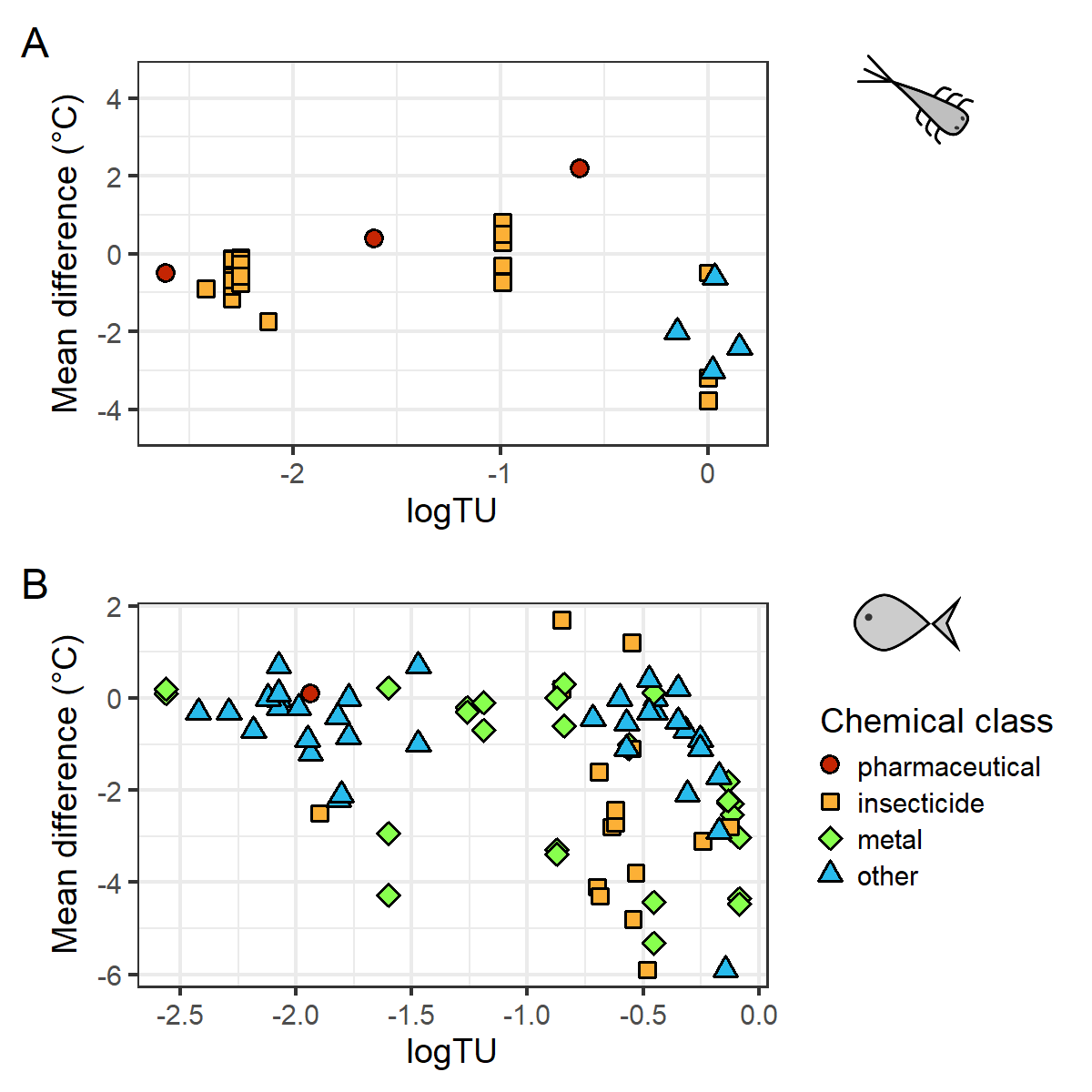

**Figure S1**. Mean differences in heat tolerance against acute chemical toxicity for (A) invertebrates (*n* = 37) and (B) fish (*n* = 80). Acute chemical toxicity is expressed as log toxic units (logTU). The dataset was trimmed to include logTU values above -3. Fill color and point shape indicates chemical class.

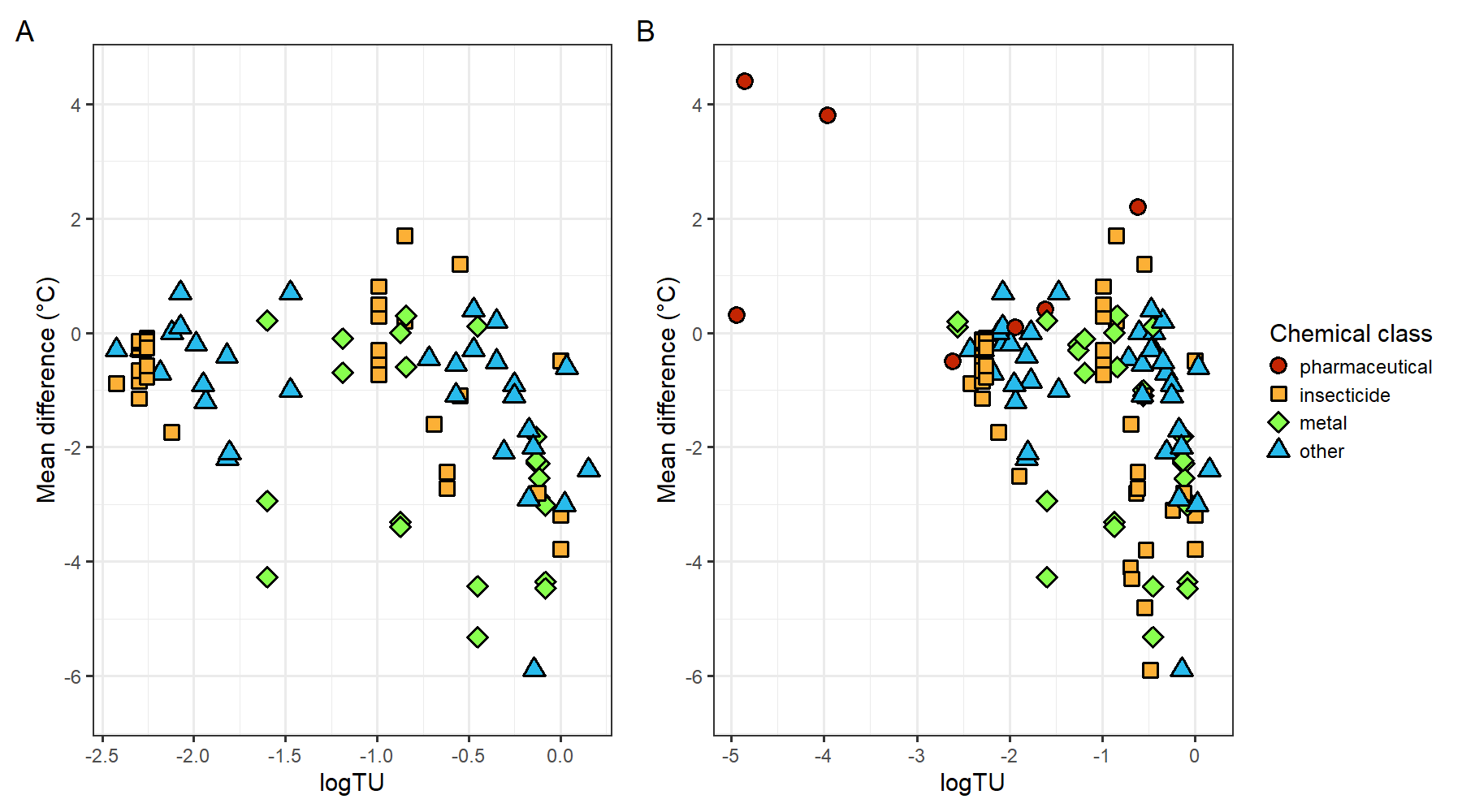

**Figure S2**. Raw data for mean difference in heat tolerance effect sizes (A) with reported measures of dispersion, and (B) untrimmed data including data without reported measures of dispersion, against the log toxic unit (logTU). The final dataset was trimmed to include only exposures tested above a logTU of -3. Fill color and point shape indicates chemical class.

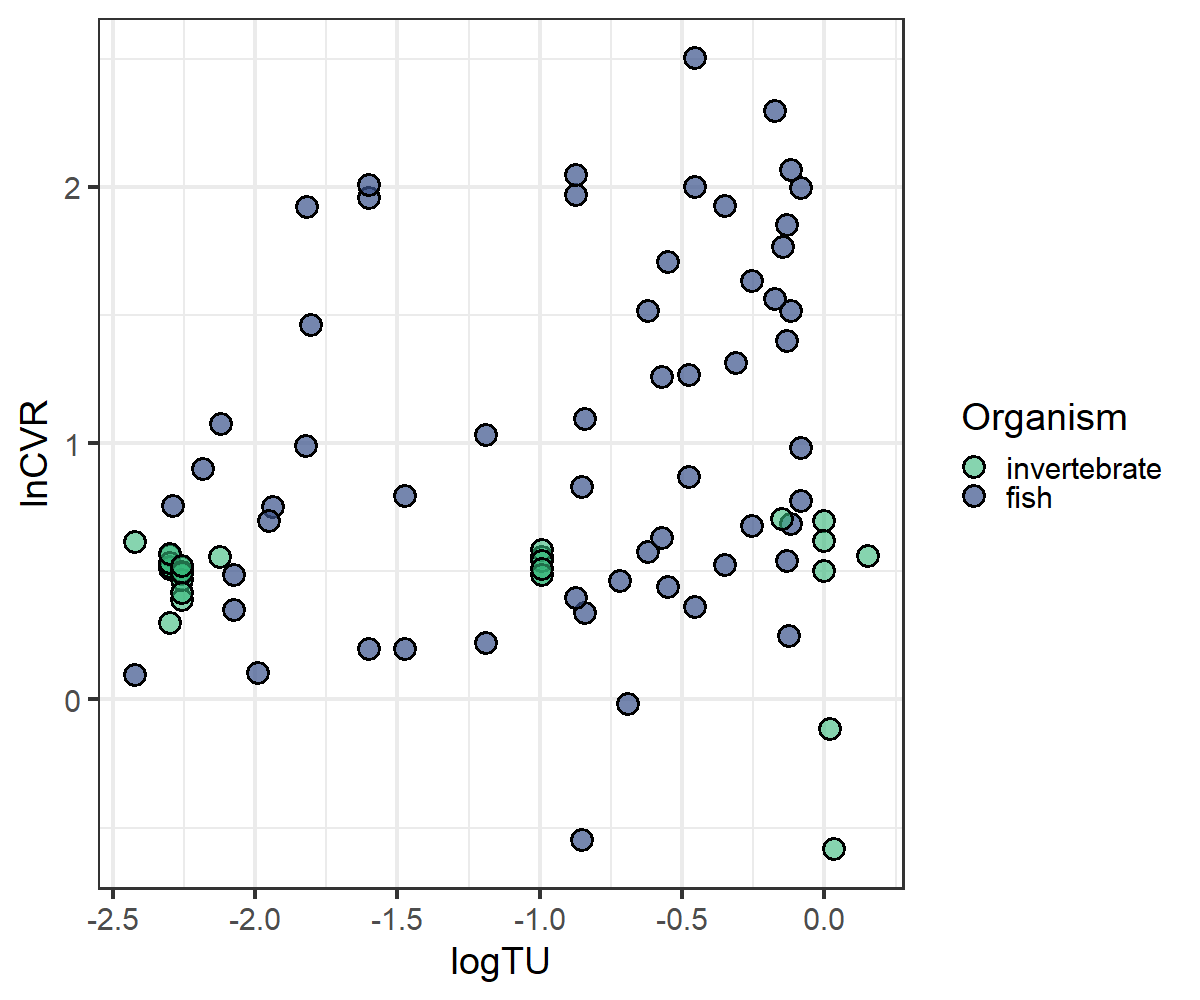

**Figure S3**. Variation against acute chemical toxicity (logTU) of each respective chemical exposure. The logarithm of the ratios of coefficients of variation (lnCVR) represents the effect size for change in variation. Fill color indicates the organism group (invertebrate or fish). A lnCVR value above zero denotes greater inter-individual variation in exposed organisms compared to control organisms, while one below zero indicates less inter-individual variation in exposed organisms compared to controls.

**Diagnostics**.

We ran diagnostic and robustness checks for all of our meta-analytic models. To check that parameters in the models were identifiable, the likelihood was profiled using the profile function in metafor (Viechtbauer, 2026). All parameters for models run with the nested structure shown in **Figure 2** were identifiable, as the likelihood profiles showed clear peaks. We also ran models with a crossed random effects structure (separate random effects for species and chemical) since both separate species and chemical random effects are plausible for our data, and analytical decisions such as the specification of fixed and random effects in statistical models can affect results (Gould *et al.*, 2025). However, the actual crossover was minimal in our dataset, since the dataset mainly consisted of unique species-chemical combinations. The models with a crossed random effects structure failed to reliably estimate parameters; profiles of the likelihood did not show a peak. To check the influence of publication year (time-lag effect), a meta-regression was run with publication year as a predictor. We checked funnel plots with effective sample size and standard error for visible asymmetry (a hallmark of publication bias) and ran a multi-level Egger regression using the square root of inverse effective sample size to check for a small-study effect indicative of publication bias.

The funnel plots with marginal and conditional residuals, plotted against the effective sample size and standard error, were largely symmetric upon visual inspection (**Figures S5-S6**). We fit an Egger regression for both the data with imputed dispersion and without; neither regression had a significant slope term (**Figure S6**). For the data without imputed dispersion, the slope of the Egger regression was close to zero (slope = -0.18, upper CI = 7.1, lower CI = -7.4, *p* = 0.96), while the data with imputed dispersions showed an upward, albeit nonsignificant, trend (slope = 4.5, upper CI = 10.6, lower CI = -1.7, *p* = 0.16). This upward trend indicates that larger, higher-powered studies (those with a smaller square root of the inverse effective sample size) tended towards more negative effects, and smaller studies tended towards effects closer to zero (**Figure S6**). This trend was only apparent in the dataset with imputed dispersion, where dispersion was not always reported in the source study. Small studies that found high variability in their estimates may have been more likely to leave variability unreported; this same variability could also hinder their ability to detect an effect. We found no significant effect of publication year, though an apparent slight upward trend (slope = 0.03, upper CI = 0.07, lower CI = -0.01, *p* = 0.11), with tests on invertebrates, insecticides, and lower acute toxicity levels slightly more prevalent after 2010 (**Figure S7**).

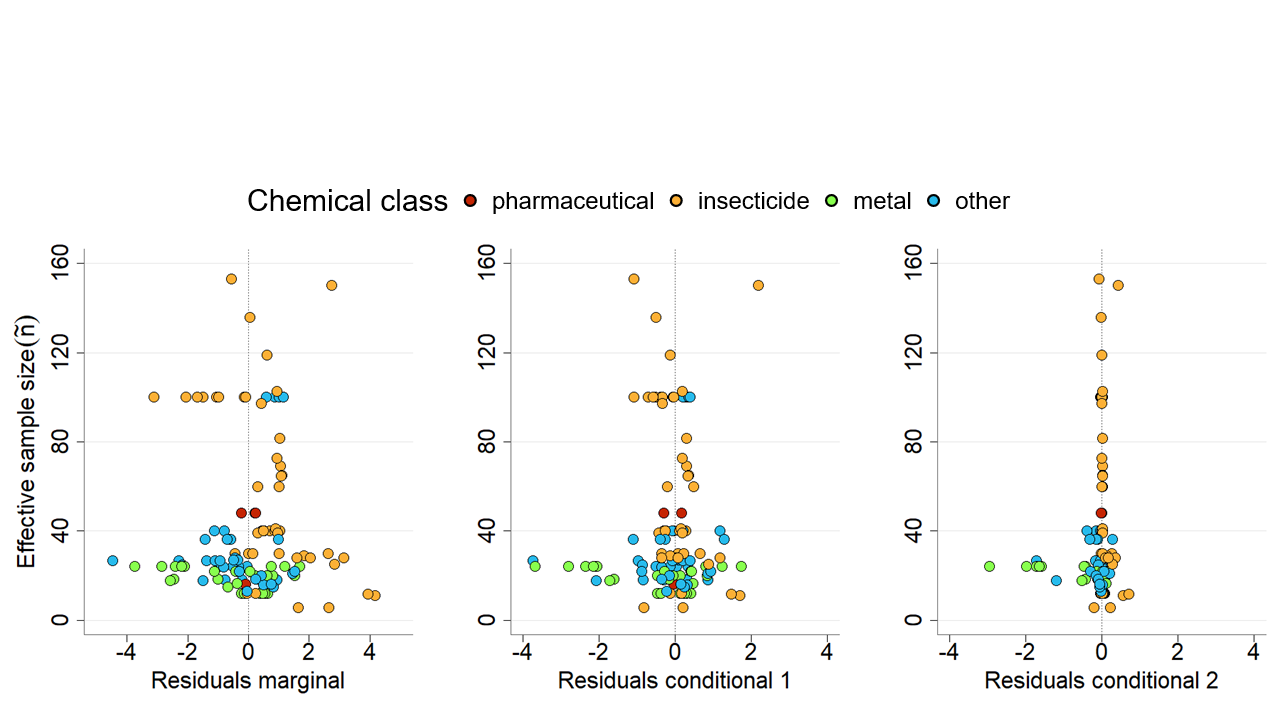

**Figure S4**. Funnel plots with marginal and conditional residuals plotted against the effective sample size of each effect size (mean differencein heat tolerance). Point color indicates chemical class.

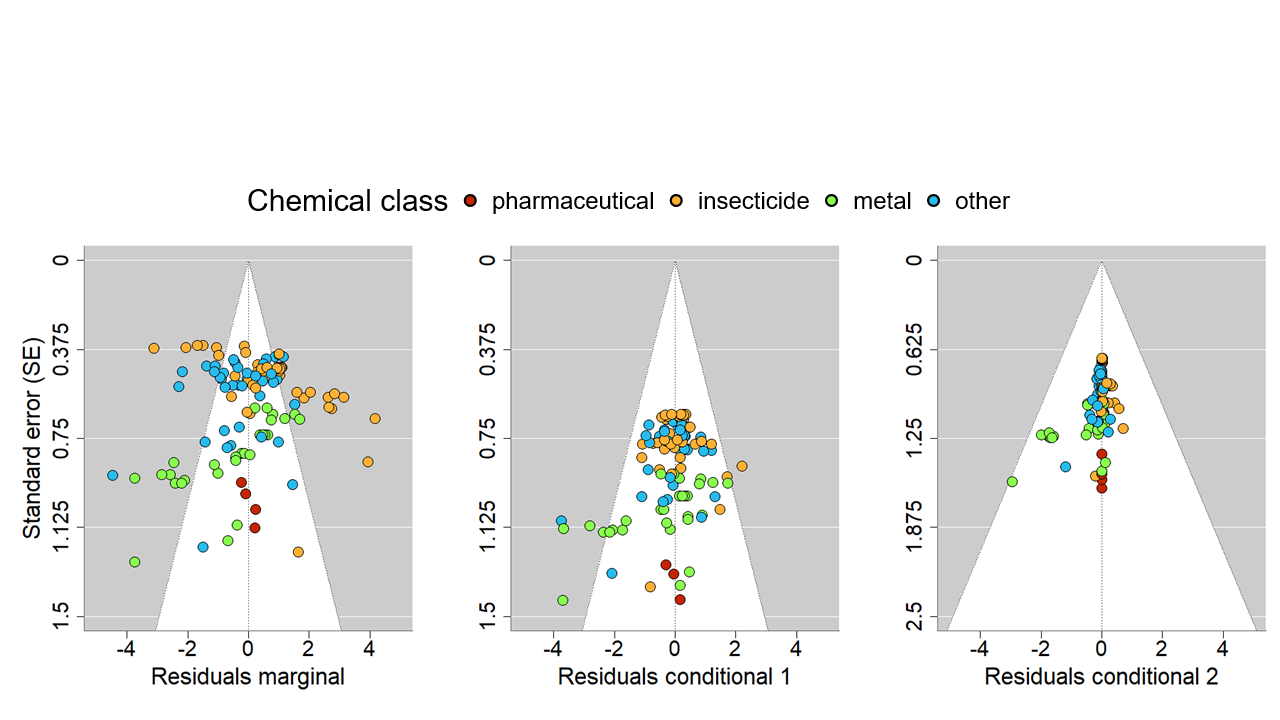

**Figure S5**. Funnel plots with marginal and conditional residuals plotted against the standard error of each effect size (mean difference in heat tolerance). Point color indicates chemical class. The points are expected to fall symmetrically around the vertical line at zero, within the area indicated by the white funnel.

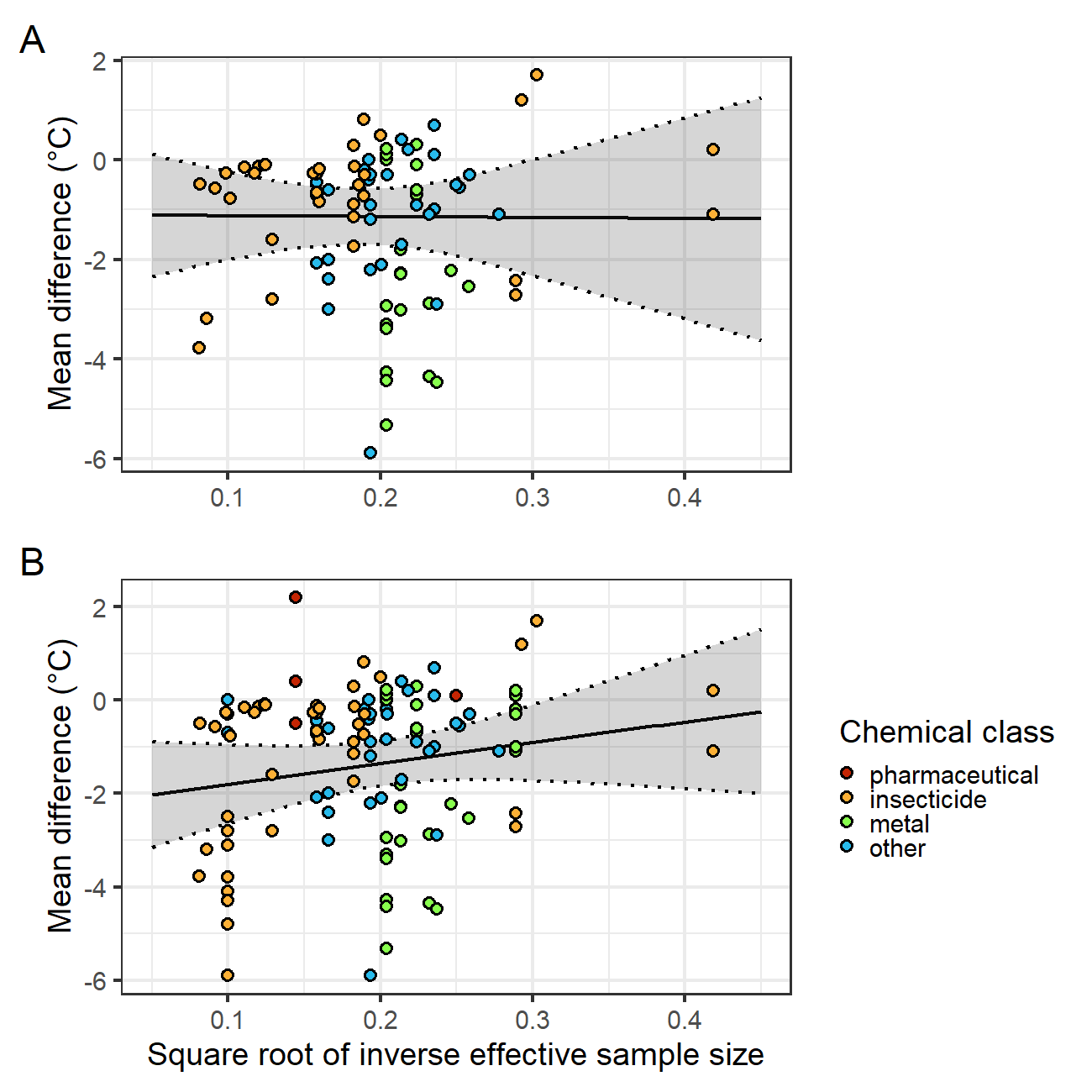

**Figure S6**. Egger regression with the square root of inverse effective sample size plotted against each effect size (mean difference in heat tolerance), for (A) data with measure of dispersion, and (B) the full dataset including imputed values. Neither regression has a significant slope, but significant intercepts indicate a small study effect. Point color indicates chemical class.

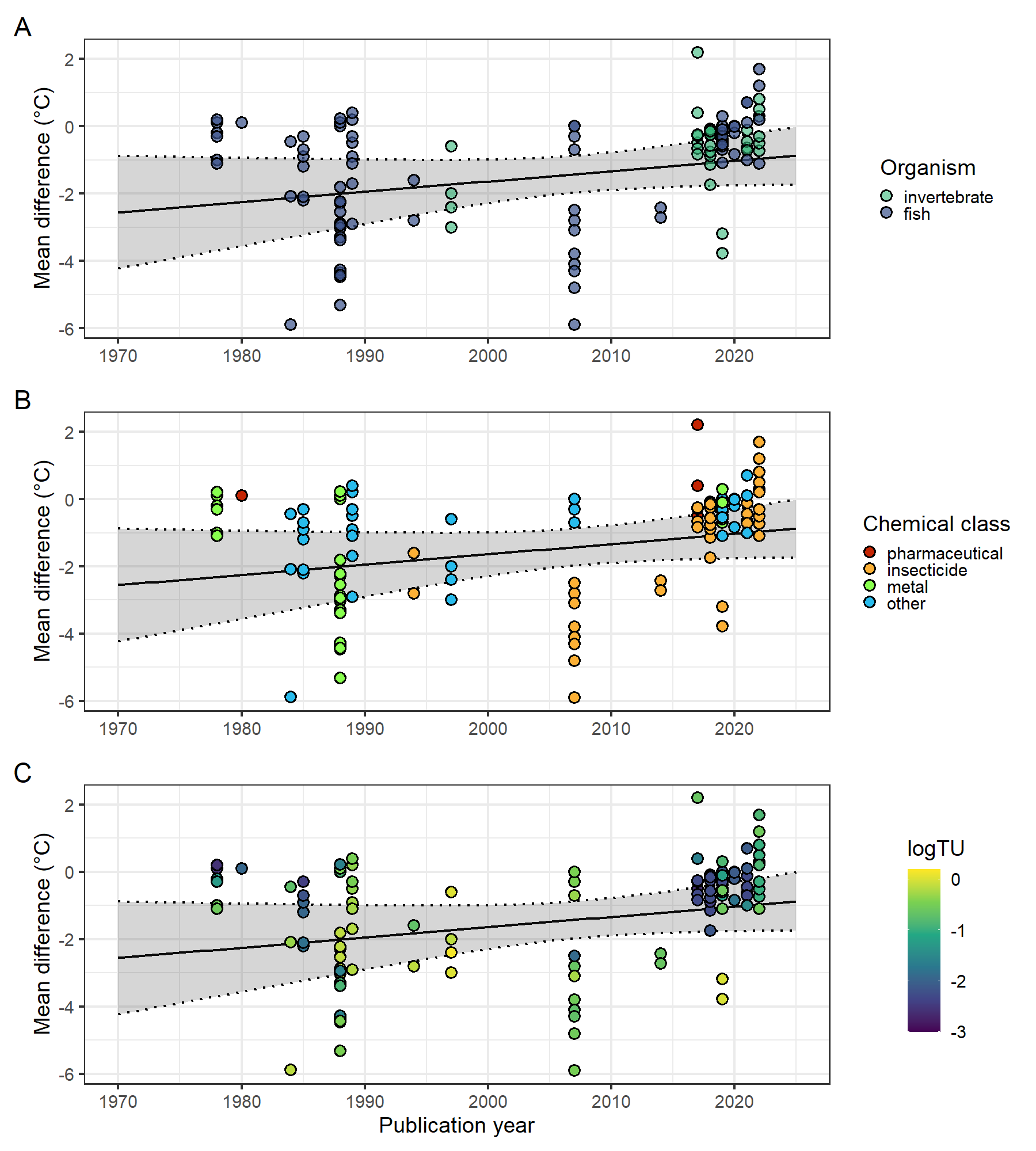

**Figure S7**. Effect sizes (mean differences in heat tolerance) plotted against publication year. Point color based on (A) organism group, (B) chemical class, and (C) acute chemical toxicity (logTU). The regression with publication year (black line with gray confidence intervals, plotted on all three panels) is not significant (*p* = 0.11). The slight, not-significant upward trend may be due to the fact that organism group, chemical class, and logTU are not evenly studied across time. Nevertheless, there is no strong support for a time-lag effect.

**References**.

Gould, E. *et al.* (2025) “Same data, different analysts: variation in effect sizes due to analytical decisions in ecology and evolutionary biology,” *BMC Biology*, 23(1), p. 35. Available at: https://doi.org/10.1186/s12915-024-02101-x.

Viechtbauer, W. (2026) “metafor: Meta-Analysis Package for R.” Available at: https://cran.r-project.org/web/packages/metafor/index.html (Accessed: April 7, 2026).
